# Cryo-electron microscopy reveals a single domain antibody with a unique binding epitope on fibroblast activation protein alpha

**DOI:** 10.1101/2024.10.18.619146

**Authors:** Zhen Xu, Akesh Sinha, Darpan N. Pandya, Nicholas J. Schnicker, Thaddeus J. Wadas

## Abstract

Fibroblast activation protein alpha (FAP) is a serine protease that is expressed at basal levels in benign tissues but is overexpressed in a variety of pathologies, including cancer. Despite this unique expression profile, designing effective diagnostic and therapeutic agents that effectively target this biomarker remain elusive. Here we report the structural characterization of the interaction between a novel single domain antibody (sdAbs), I3, and FAP using cryo-electron microscopy. The reconstructions were determined to a resolution of 2.7 Å and contained two distinct populations; one I3 bound and two I3 molecules bound to the FAP dimer. In both cases, the sdAbs bound a unique epitope that was distinct from the active site of the enzyme. Furthermore, this report describes the rational mutation of specific residues within the complementarity determining region 3 (CDR3) loop to enhance affinity and selectivity of the I3 molecule for FAP. This report represents the first sdAb-FAP structure to be described in the literature.

## Introduction

Single domain antibodies (sdAbs or VHH), which were first detected in the sera of *Camelidae*, are a class of immunoglobulins that lack light chains and consist of only one heavy chain with a single variable domain^1–3^. When compared to conventional heavy chains (VH) of regular IgG molecules, their three complementarity determining regions (CDRs) are enlarged to provide a greater surface area for antigen interactions making them well-suited for binding restricted sites such as cavities or sterically hindered epitopes. Moreover, they contain additional hydrophilic amino acids within the conserved framework region. VHH proteins retain high affinity and specificity for their target antigens, with low off-target accumulation. Compared to the stability exhibited by a conventional antibody, they are unexpectedly robust due to their high refolding capacity, recovering from chemical denaturation with minimal damage to functionality. Furthermore, unlike conventional antibodies, they can tolerate environmental conditions associated with radiochemistry including high temperatures, elevated pressures and non-physiological pHs. Additionally, nanobodies are relatively simple and inexpensive to produce on the milligram scale in a laboratory setting since they lack post-translational modifications and can be synthesized in microbial systems. As a result, the last three decades have witnessed explosive growth in research related to these molecules to use them as diagnostic and therapeutic agents for a variety of pathologies. As of 2020, there were over 15 clinical trials involving sdAbs^4,5^.

The dipeptidyl peptidase (DPP) family of proteins are metalloproteases that cleave the N-terminal dipeptide from peptides with Pro or Ala in the penultimate position; the family’s substrates include growth factors, chemokines, neuropeptides, vasoactive peptides, and extracellular matrix molecules such as collagen. This family has seven family members including DPP4, DPP8, DPP9, DPPII, prolyl carboxypeptidase (PRCP), prolyl oligopeptidase (PREP) and fibroblast activation protein alpha (FAP or seprase)^6,7^. Of these family members significant research activity has revolved around FAP, which is a 170 kDa type II transmembrane serine protease since it is unique among this enzyme family because of its endopeptidase activity and substrate selectivity^8–10^. Moreover, unlike other members of this protein family, FAP exhibits a unique expression profile and is considered a robust biomarker of pathology since its demonstrates negligible expression in normal adult tissues, but is prominently expressed in a variety of pathologies including cancer, arthritis, atherosclerosis and fibrosis. Several reports describe strategies to target FAP expression for imaging and therapy using peptides, antibodies, antibody fragments, nanoparticles and small molecules have appeared in the literature^11–18^. Recently, single domain antibodies targeting FAP have been described as potential theranostic agents. For example, Xu et al. identified two novel anti-FAP VHH proteins that were engineered to contain the Fc fragment of IgG4^19^. These recombinant proteins were radiolabeled with zirconium-89 (^89^Zr: t_½_ = 78.4 h, β^+^: 22.8 %, E_β+max_ = 901 keV; EC: 77%, E_γ_ = 909 keV) and lutetium-177 (^177^Lu^3+^: β^−^ - emitter: t_1/2_ = 6.7d; E ^−^ = 0.497 MeV)^20,21^. Ex vivo biodistribution analysis of the ^89^Zr-agent revealed good tumor uptake at later time points, while therapy studies with the ^177^Lu-agent demonstrated tumor growth control without significant animal toxicity. Recently, a study by Dekempeneer, et al. revealed additional single domain anti-FAP antibodies with K_D_ values in the nano-to-picomolar range. These VHH molecules were radiolabeled with several PET, SPECT and therapeutic radioisotopes^20,22,23^. These agents exhibited specific accumulation in human FAP^+^ tumors, while being excreted rapidly in most cases. The proteins radiolabeled with actinium-225 (^225^Ac^3+^: α^++^ - emitter: t_1/2_ = 10 d; E_αmax_ = 6-8 MeV) demonstrated kidney retention but provided tumor growth control in FAP^+^ tumor bearing mice. Collectively, these publications demonstrate the potential of anti-FAP single domain antibodies in the development of theranostics. However, despite interesting data reported, neither group of authors specifically described the binding epitope of the reported VHH molecules. Unfortunately, this lack of structural information hinders further development from a rational drug design perspective.

Cryo-electron microscopy (cryo-EM) is a biophysical technique that enables the structural determination of large and/or dynamic macromolecules^24–32^. As a complimentary technique to X-ray crystallography and NMR, cryo-EM has become an important tool in the drug discovery process and a valuable asset for structural biologists who wish to interrogate the structure and function of large protein complexes at near atomic resolution^33^. While still evolving, cryo-EM methodologies have become robust enough to confidently model amino acid side chains and ligands into the density maps. These improvements are continuing to transform the drug discovery process and has helped in faster lead molecule identification. This report describes the utilization of cryo-EM, mass photometry (MP) and biolayer interferometry (BLI) to characterize the interaction of the single domain antibody, I3 with the serine protease, fibroblast activation protein alpha. To our knowledge, this is the first report to describe the binding interaction between FAP and an anti-FAP VHH using cryo-EM.

## Results

### Characterization of I3 binding to FAP

To analyze the complex and determine the stoichiometry of I3 binding to FAP, mass photometry (MP) was employed. The FAP protein was confirmed to be a dimer (198 kDa) at the low nM concentrations used for mass photometry (Fig. 1A), matching the previously determined structure^34^. Characterization of affinity and binding kinetics were done using biolayer interferometry (BLI). A site-specific biotinylated version of FAP and an MBP-I3 construct were used for BLI. The MBP-I3 construct was chosen for its larger size, and therefore greater response in BLI. The biotinylated FAP was immobilized on streptavidin coated biosensors and dipped into a range of MBP-I3 concentrations to measure association and then into buffer wells to measure dissociation (Fig. 1B). The affinity (K_d_) was determined to be 2.0 ± 0.3 μM by fitting to the kinetic data and 2.7 ± 0.1 μM by steady state analysis (Fig. S1). Additionally, the kinetic rates for association (k_on_) and dissociation (k_off_) were 1.8 (± 0.3) *10^5^ M^−1^s^−1^ and 3.6 (± 0.2) *10^−1^ s^−1^, respectively. Detection of the FAP complex with SUMO-I3 by MP required a cross-linking reagent due to the weak affinity and fast dissociation rate (see methods). Complex formation was confirmed and contained a heterogenous mix of populations including FAP alone, FAP + SUMO-I3, and FAP + 2 SUMO-I3 (Fig. 1C). This verified 1 I3 binding site per FAP monomer.

**Figure 1.**
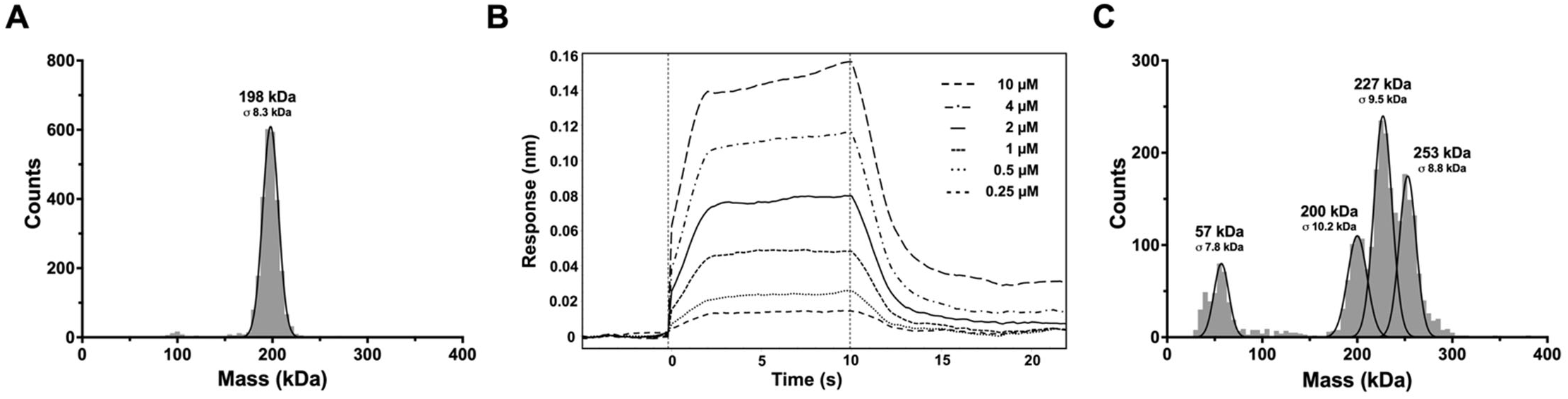
Mass photometry analysis of FAP with SUMO-I3 complexes. (A) Mass distribution of 15 nM FAP alone. The molecular weight (MW) observed by MP for FAP is 198 ± 8.3 kDa, which agrees well with the predicted MW of the dimeric FAP (170 kDa) with glycosylation. (B) Sensorgram from bio-layer interferometry data showing binding of FAP with increasing concentrations of MBP-I3. Data is representative of triplicate measurements. (C) Mass distribution of 15 nM BS3 cross-linked FAP with SUMO-I3 in 1:5 molar ratio. The MWs observed are 57 ± 7.8 kDa, 200 ± 10.2 kDa, 227 ± 9.5 kDa, and 253 ± 8.8 kDa, which corresponds to the expected MWs of two SUMO-I3 (54 kDa), glycosylated FAP alone (198 kDa, panel 1A), and FAP with one SUMO-I3 (225 kDa) or 2 SUMO-I3 (252 kDa) molecules bound, respectively.

### FAP-I3 complex structure

To determine the epitope for I3, the structure of SUMO-I3 bound to FAP was determined by cryo-EM. Structure determination used cross-linked sample as it resulted in far greater complex compared to uncross-linked sample (data not shown). Like the dual populations observed in the MP data, particles with both one and two I3 molecules bound to FAP were isolated during the cryo-EM data processing workflow (Fig. S2, Fig S3 and Table 1). The reconstructions for both FAP I3 complexes were determined to 2.7 Å for one I3 bound in C1 symmetry and two I3 molecules bound in C2 symmetry (Fig. S2). Local resolution maps and final structural models for each complex are shown in Fig. 2. Due to the flexible portion between the SUMO and I3 regions of the fusion protein, the SUMO region is not observed in the reconstruction. The local resolution throughout most of the FAP core region is ∼2.6 Å and ∼3.0 Å at the FAP-I3 interface. Both FAP molecules and the I3 molecules from each complex overlay well with very subtle differences (C-alpha RMSDs, FAP dimer 0.141, I3, 0.199). I3 interacts with FAP through its CDR3 loop and FR2 region, instead of the more typical interaction with all three CDR loops (Fig. 3A). Specific residues involved in the FAP-I3 interaction can be seen in Fig. 3B. FAP Y274 has multiple interactions with I3 and sits within a pocket formed by I3 (Fig. 3B,C). The electrostatic surface map of I3 shows that the one loop from FAP engages a relatively uncharged region while the second FAP loop interacts with a positively charged region (Fig. 3C). The epitope footprint on the surface of FAP is shown in Fig. 3D. The following FAP:I3 residue pairs have a hydrogen bond interaction; Y274:P108, Y274:W47, E325:S109, and D326:V107. Additionally, there is a π-stacking interaction between Y271:F110.

**Table 1.**
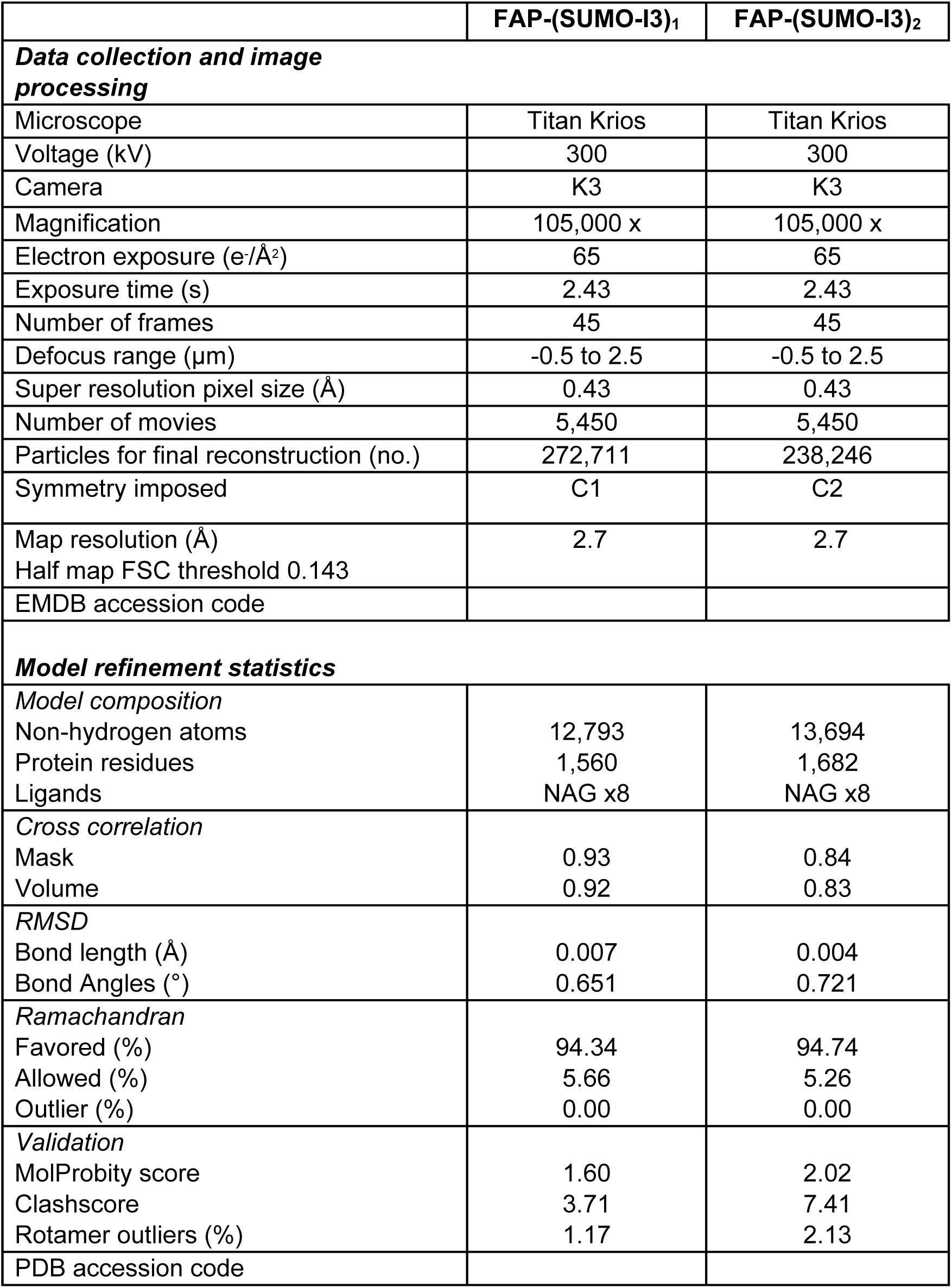
Cryo-EM data collection parameters and model refinement statistics.

**Figure 2.**
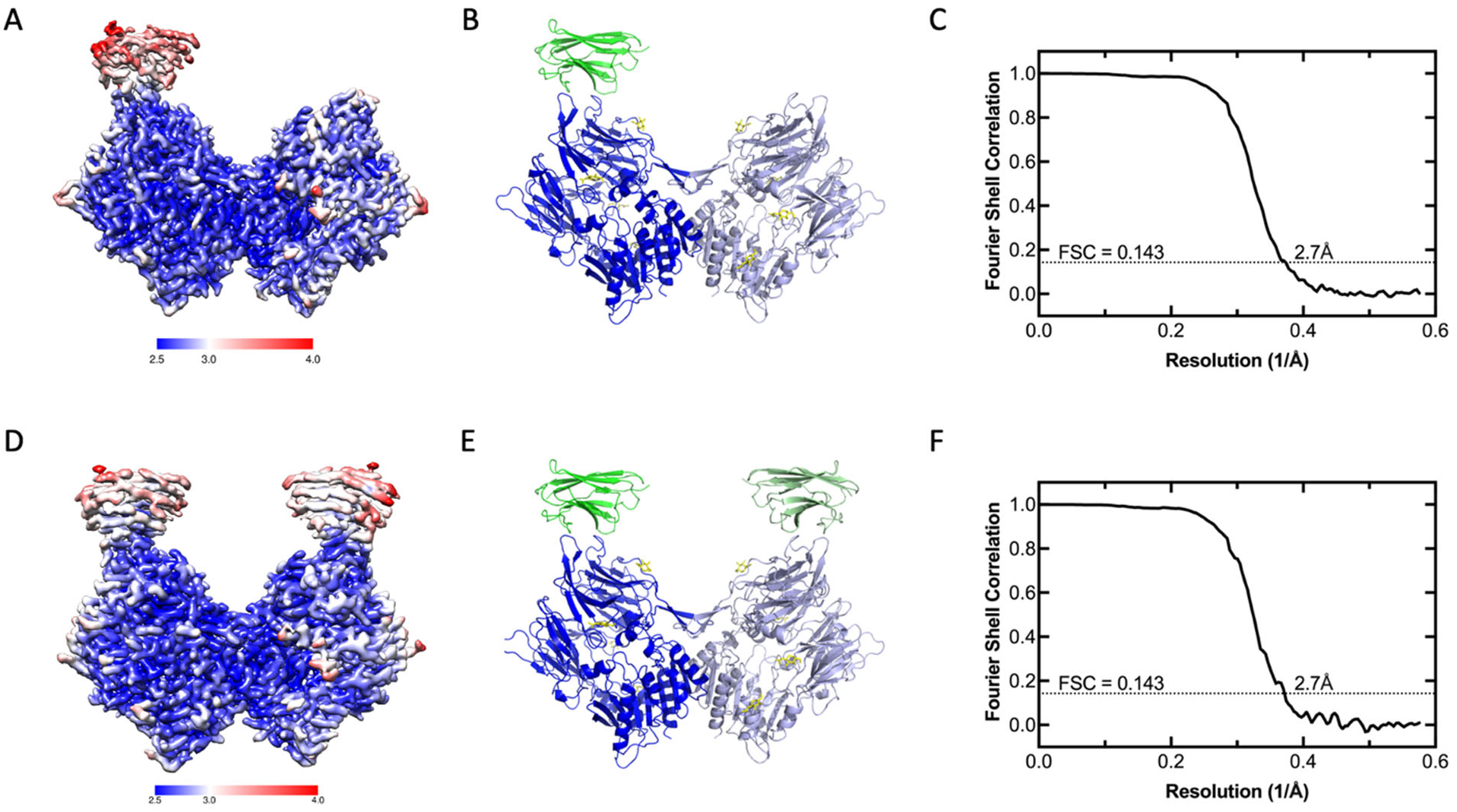
Cryo-EM structures of FAP-I3 complexes. (A) FAP-I3 reconstruction with local resolution map. (B) FAP-I3 model. (C) FSC curve for the 3D reconstruction of the cryo-EM map of FAP-I3. The average resolution is estimated to be 2.7 Å based on the FSC value of 0.143. (D) FAP-(I3)_2_ reconstruction with local resolution map. (E) FAP-(I3)_2_ model. (F) FSC curve for the 3D reconstruction of the cryo-EM map of FAP-(I3)_2_. The average resolution is estimated to be 2.7 Å based on the FSC value of 0.143. Cartoon models in (B) and (E) show the FAP dimer in two shades of blue and I3 in green. Yellow sticks represent sites of glycosylation on FAP.

**Figure 3.**
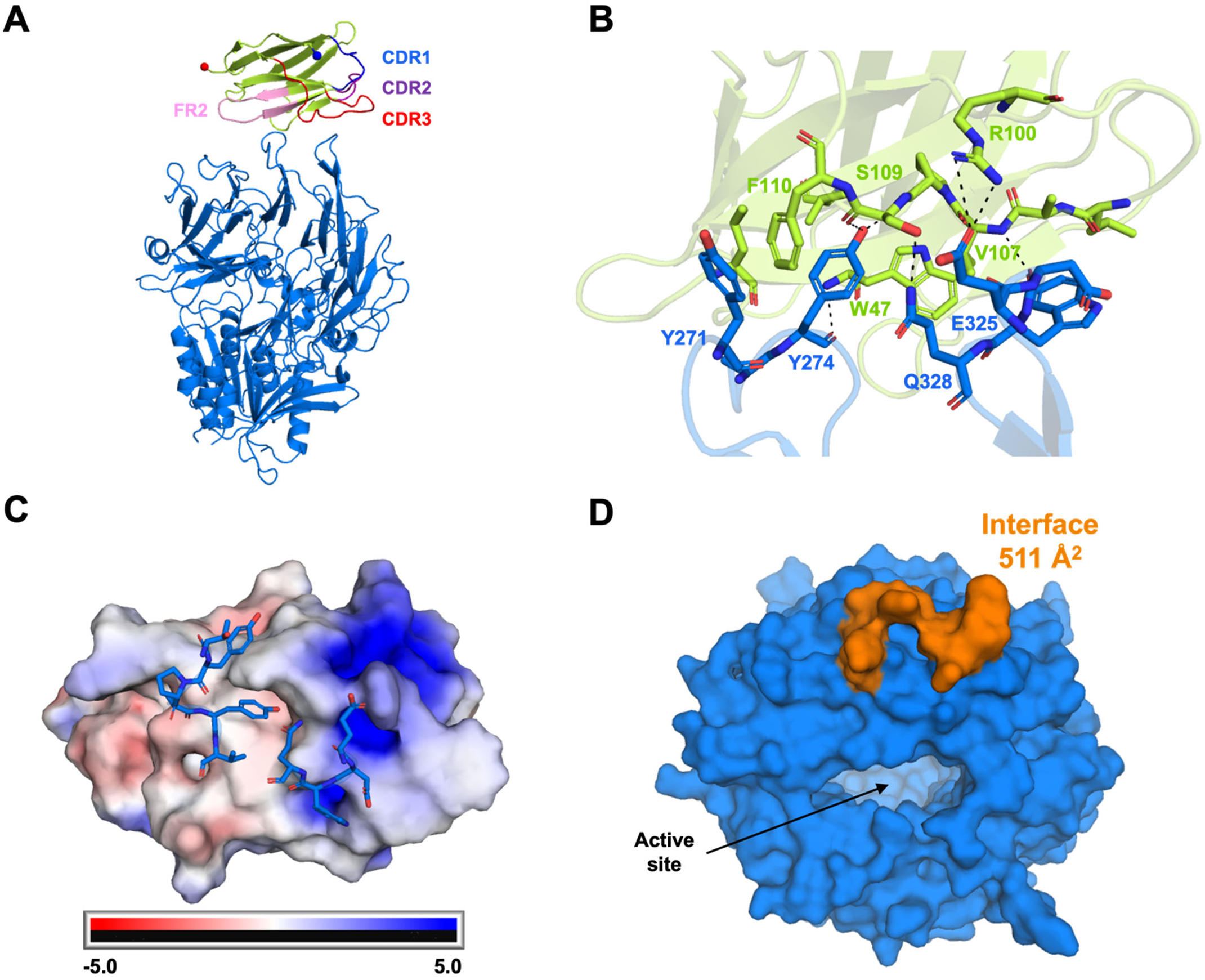
Interactions of FAP with I3. (A) Overview of I3 bound to FAP showing various VHH regions typically involved in the paratope. Blue and red spheres correspond to the first and last residues modeled for I3. (B) Residues involved in specific interactions at the FAP-I3 interface. (C) An electrostatic surface map of I3 and residues from FAP involved in epitope formation in blue sticks. (D) A surface representation of FAP highlighting the I3 epitope region and its location relative to the active site.

### In silico rational design to enhance I3

Since the affinity of I3 for FAP was weak compared to typical sdAbs, rational mutations were chosen to improve the affinity. The sites V107 and S109 were identified on I3 that could potentially benefit from having a mutation with positive electrostatic potential or an aromatic residue (Fig. 4A). In silico mutations were chosen for V107 and S109 that could potentially increase and decrease (as a control) the FAP-I3 interaction (Fig. 4B, C). The calculated changes in affinity (dAffinity) and stability (dStability) from the original sequence for both sites showed approximate trends expected with various mutations. The Arg mutations were projected to provide the largest increase in both affinity and stability. The comparison of hydrogen bonds, salt bridges, and π-stacking interactions at the interfaces shows S109R to form 2 new hydrogen bonds and a salt bridge, which positively increases affinity and stability (Fig. 4D).

**Figure 4.**
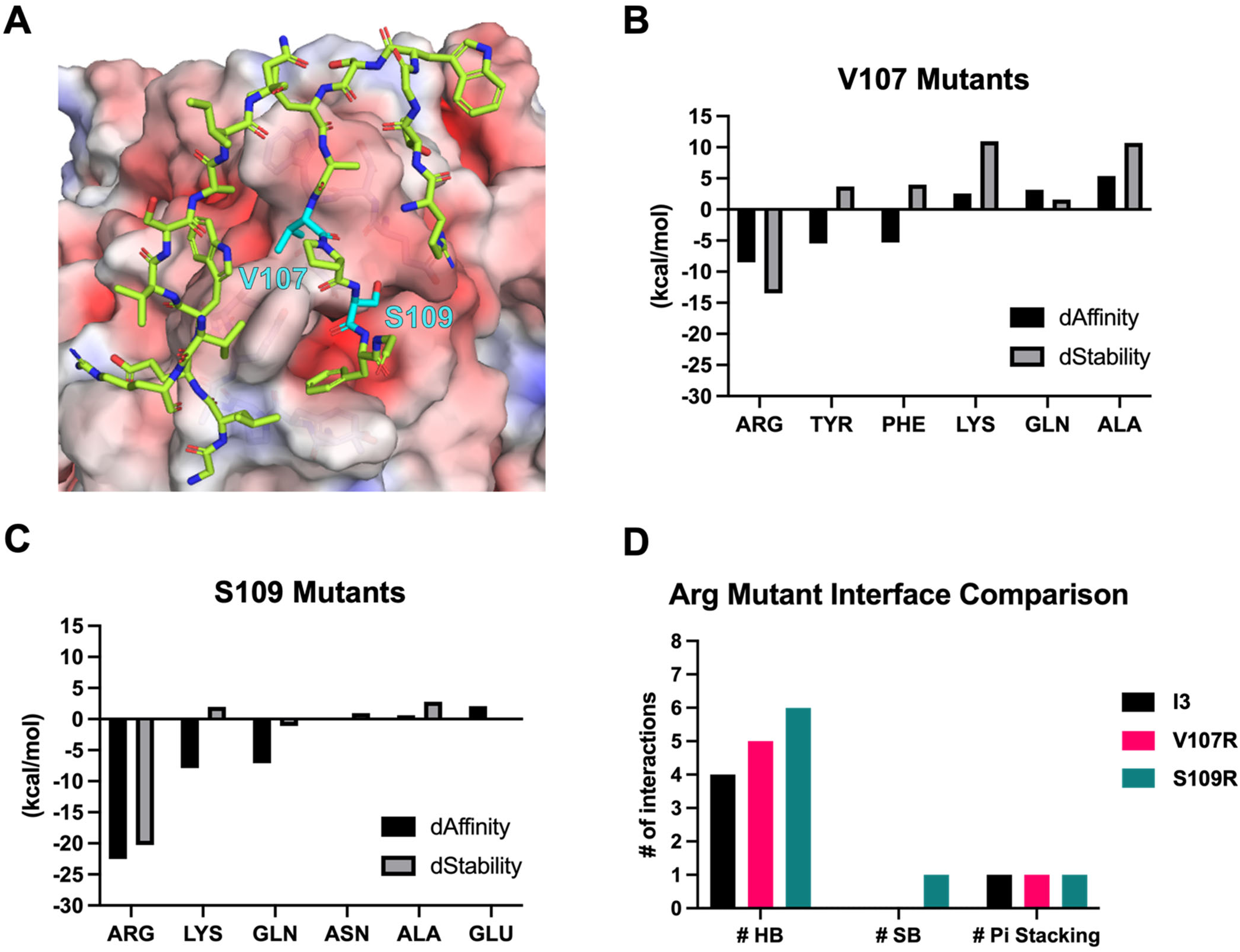
In silico affinity maturation of I3. (A) An electrostatic surface map of FAP and some residues of I3 near the surface in green sticks. Residues chosen for mutation are in cyan. (B) V107 mutation change in affinity and stability results compared with the original I3 sequence. (C) S109 mutation change in affinity and stability results compared with the original I3 sequence. (D) Comparison for the number of hydrogen bonds (HB), salt bridges (SB), and pi stacking interactions present at the interfaces of I3 and the V107R and S109R mutants.

### Comparison of FAP epitope to homologous DPP4 protein

FAP belongs to the dipeptidyl peptidase (DPP) family and shares 52% sequence identity (71% similarity) with DPP4. Both FAP and DPP4 are dimeric and share high overall structural homology (Fig. 5A). Comparison of the FAP I3 epitope region to the same region in DPP4 shows distinct differences in orientation of the crucial FAP loop containing Y274 (Fig. 5B). The equivalent loop in DPP4 (residues 275-283) protrudes less from the overall protein. Analysis of the sequences from this epitope region reveals large differences in residue composition in addition to the structural conformation (Fig. 5C). The lack of MBP-I3 binding to DPP4 was tested and confirmed by BLI (Fig. 5D).

**Figure 5.**
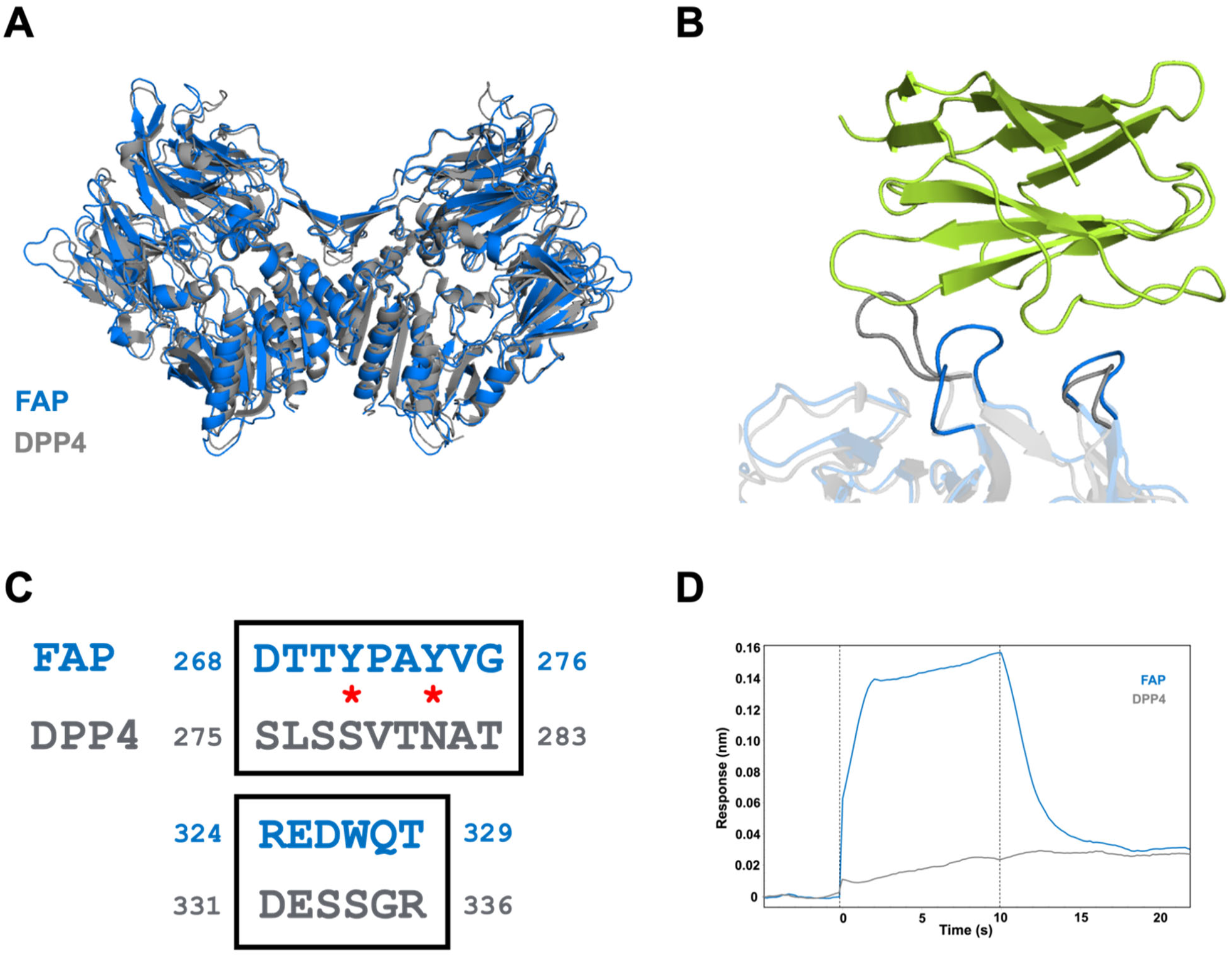
FAP epitope comparison with DPP4. (A) Overall structure-based alignment of FAP with DPP4. (B) Highlight of FAP epitope region comparison shows conformational differences in DPP4. (C) Structure based sequence alignment of the FAP epitope region with DPP4. Two key residue differences are indicated by red asterisks. (D) BLI binding data showing MBP-I3 (10 μM) specifically interacts with FAP and not DPP4.

## Discussion

The ability to identify an antibody binding region or epitope of a protein considered to be an important biomarker of disease has important implications for disease diagnosis, vaccine development, and elucidating disease mechanisms^35–39^. Additionally, the characterization of an antigen binding region enables the characterization of therapeutic antibodies and has important intellectual property implications. Epitope mapping is the determination of which amino acid sequences and three-dimensional interactions directly contribute to the affinity between an antibody or its derivatives and a specific antigen. Furthermore, epitope mapping allows investigators to study how the binding of specific epitopes may alter protein function.

The present study demonstrates that the single domain antibody I3, which has not been previously disclosed within the Structural Antibody Database (SAbDab), binds to the extracellular surface of FAP by making important contacts through its CDR3 at a unique epitope that is distinct from the active site. These unexpected results may have a profound effect on how this molecule may be utilized for diagnostic or therapeutic purposes since the role of FAP has been shown to be context and disease dependent^9,10,40^; the expression of FAP may be beneficial under some circumstances such as pulmonary fibrosis while detrimental in other circumstances such as cancer^41^. While antigen binding without enzyme inhibition still needs to be confirmed, being able to target the protein without inhibiting function may be a viable path to the successful development of a new class of diagnostic and therapeutic molecules for a variety of disease states.

Also, given the current data supporting the cell-surface heteroprotein complex formation that occurs between FAP, dipeptidyl peptidase IV (DDP4), matrix metalloproteinases (MMPs), integrins and uPAR, understanding how I3 binds to FAP may facilitate the rational design of bi-specific ligands that target FAP and additional partners at the cell surface, and offer another strategy for improved development of agents with diagnostic or therapeutic value^42^. A structural alignment comparison of FAP with DPP4, DPP8, and DPP9 shows the I3 footprint region differs for these S9 family members (Fig. S4). This is an important factor of the I3 epitope as these proteins all share high sequence and structural similarity. Finally, since sdAbs are often used to stabilize larger proteins, the use of I3 may have utility as a molecular chaperone when studying FAP and unexplored aspects of its biology that cannot be elucidated using current structural and molecular biology techniques^43^. Such areas include the true expression pattern of FAP on different cell types, more rigorously distinguishing the enzymatic and non-enzymatic roles of FAP in healthy and diseased tissues and understanding the mechano-signaling aspects of FAP.

While this work reveals a previously unknown and unique binding epitope on the FAP protein, several limitations of this work should be considered. Foremost, the binding affinity of I3 for FAP is low when compared to other sdAbs reported previously^19,22^. While disappointing, this is not surprising since I3 was derived from a naive camelid library in contrast to other sdAbs that were derived from immunized animals. While such targeted libraries can yield several molecules with high affinity for their target protein, the cost and logistics associated with identifying these molecules is extremely high and complicated, respectively. Thus, these development strategies are available only to commercial entities with the capital and organizational infrastructure to conduct such lengthy discovery experiments. Furthermore, while our *in silico* design has predicted new variants that will achieve better affinity and selectivity that I3, these variants have not been evaluated nor have their structural interactions with FAP been elucidated to date. However, these experiments are currently underway in our laboratories and will be communicated in subsequent publications.

Finally, this work represents a powerful example of how cryo-EM can be applied to the drug discovery process and the rational design of sdAbs. Furthermore, it is expected to play an ever more important role now that the *de novo* design of sdAbs is on the horizon. Recently, Bennett et al. described a computational strategy to accurately design sdAbs and tested this methodology by creating sdAbs specific for influenza hemagglutinin ^44^. In their report the authors demonstrated that the computationally derived sdAbs were nearly identical in CDR conformation and overall binding to the binding models developed using conventional molecular biology and elucidated using cryo-EM. Thus, as these techniques become more sophisticated and routine due to technological advancements in machine learning, the more tedious and time-consuming way of generating sdAbs, which involve animal immunization and library screening, will become secondary options; these artificial intelligence (AI) tools are expected to form the foundation of a new period in the rational design of antibodies and their derivatives.

In conclusion, to our knowledge, this work is the first report describing a unique epitope on the FAP protein that is occupied by the single domain antibody, I3. This work also reveals the important residues necessary for the FAP-I3 interaction and rationalizes why I3 selectively binds to FAP and not its closest S9 family member DPP4. Considering the current research interest in FAP as a therapeutic target in a variety of disorders including cancer, fibrosis, arthritis and cardiovascular disease, this work will assist investigators in developing theranostic agents to assess mitigate these disease states.

## Methods

### Reagents and equipment

Unless noted, chemicals and materials were purchased from Sigma-Aldrich Chemical Co. (St. Louis, MO, USA) or ThermoFisher Scientific, Inc. Solutions were prepared using ultrapure water (18 MΩ-cm resistivity). Protein purification was accomplished using an NGC FPLC system (Bio-Rad, Hercules, CA). Human recombinant fibroblast activation protein alpha and dipeptidyl peptidase IV (DPP4) were purchased from Biolegend (San Diego, CA). The single domain antibody, I3 was isolated from a naïve camelid library and purchased from Neoclone Biotechnologies, International (Madison, WI). The sequence coding for I3, was synthesized at Integrated DNA Technologies (Coralville, IA).

### Preparation of the single domain antibody, I3

The coding sequence of I3, along with an N-terminal FLAG tag, was cloned in the pRham^TM^ N-His SUMO Kan vector (Lucigen) following manufacturers’ recommendations. This expression construct facilitated the expression of I3 fused to an N-terminal 6X His-SUMO-FLAG tag, which is referred as SUMO-I3. The SUMO-I3 construct was transformed into Shuffle T7 *Escherichia coli* strain (New England Biolabs) for expression, which allows for the cytoplasmic disulfide bond formation. The transformed cells were selected on LB agar plates containing 30 µg/ml kanamycin and cells were grown at 30°C. A single colony was then inoculated in 25 ml LB media supplemented with 30 µg/ml kanamycin, 0.2% (w/v) rhamnose and 0.075% (w/v) glucose and the culture was grown at 30°C at 220 rpm for overnight. The cells were harvested by spinning them at 4,000 rpm for 10 minutes at room temperature. The cell pellet was resuspended in 4 ml of Tris buffered saline (TBS; 50 mM Tris, 150 mM NaCl pH 8) containing 0.1 mg/ml lysozyme. The cell suspension was incubated on ice for 20 minutes and cells were lysed by sonicating them at 70% amplitude with twenty pulses of 10 seconds each followed by a 40 second gap. The cell lysate was centrifuged at 13,000 rpm at 4°C and the supernatant was collected. The supernatant was then incubated with 500 μl Ni-NTA resin, pre-equilibrated in TBS, for 30 minutes at 4°C. The resin bound to the recombinant sdAb was then packed in a Poly-Prep chromatography column (Bio-Rad, Hercules, CA, USA) and washed with 10 column volumes of TBS followed by washing with 10 column volumes each of TBS containing 10 mM and 20 mM imidazole. Stepwise elution of the his-tagged sdAb was performed with two column volumes each of TBS containing 50-, 100-, 200- and 300-mM imidazole. Chromatography was done in a gravitational flow mode with the flow rate of ∼0.5 ml/min. Fractions were analyzed for the presence of the sdAb on 12% SDS-PAGE and visualized after Coomassie blue staining. The major fraction was subjected to size exclusion chromatography using a Superdex 75 column (Cytiva) on an NGC FPLC system (Bio-Rad) in 100 mM PBS pH 7.2 buffer. The peak fractions were collected and concentrated using an Amicon ultra centrifugal filter with a 3 kDa MWCO (MilliporeSigma).

The coding sequence of I3 was also cloned into a custom pMAL-c5X vector (New England Biolabs) which allows for sdAb expression with an N-terminal MBP-fusion protein cleavable with TEV protease and a C-terminal 6X His-tag. The protein was expressed in Shuffle T7 *Escherichia coli* strain (New England Biolabs) for expression. The transformed cells were selected on LB agar plates containing 100 µg/ml ampicillin and cells were grown at 30°C. A single colony was then inoculated in 100 ml LB media supplemented with 100 µg/ml ampicillin and the culture was grown at 30°C at 220 rpm for overnight. Two flasks containing 1 L LB media each containing 100 ug/mL ampicillin was inoculated with 10 mL of overnight grown culture and the culture was grown at 30°C till OD600 reached 0.5-0.6. The culture was then induced by adding 0.5 mM IPTG and the culture was grown overnight at 18°C. The cells were harvested at 5,000 rpm for 30 minutes at 4 °C. The cell pellet was resuspended in buffer (50 mM Na-phosphate, 300 mM NaCl, 10% glycerol and 5 mM Imidazole, pH 7.5) with EDTA-free protease inhibitor cocktail (Roche), 0.1 mg/mL lysozyme and DNase I. The cell suspension was incubated on ice for 20 minutes and cells were lysed by sonication. The cell lysate was centrifuged at 35,000 rpm for 60 minutes at 4°C and the supernatant was collected. The protein was purified with a 5mL Ni-NTA column using a 50 mL gradient of 5 to 300 mM imidazole elution. The pooled elution fraction was subjected to size exclusion chromatography on a Superdex 200 column (Cytiva) using an NGC FPLC system (Bio-Rad) in 1x Dulbecco’s phosphate-buffered saline (DPBS) pH 7.2 buffer. The peak fractions were collected and concentrated using an Amicon ultra centrifugal filter with a 3 kDa MWCO (MilliporeSigma). The sdAbs were quantitated by measuring their absorbance at 280 nm on a nanodrop instrument (Thermo Scientific). The proteins were stored on ice for subsequent experiments.

### Cross-linking of FAP with SUMO-I3

Human FAP (2 µM, Biolegend) was mixed with 5-fold molar excess of SUMO-I3 sdAb in the binding buffer (100 mM PBS pH 7.2) and incubated at room temperature overnight. The complexes were then incubated with 1 mM bis(sulfosuccinimidyl)suberate (BS3) cross-linker (Pierce) for 30 minutes at room temperature and then the free cross-linker was quenched by adding 50 mM Tris-HCl pH 8 buffer.

### Mass Photometry

Mass photometry experiments were performed on a Refeyn TwoMP mass photometer (Refeyn Ltd, Oxford, UK). Microscope coverslips (24 mm x 50 mm, Thorlabs Inc.) were cleaned by serial rinsing with Milli-Q water and HPLC-grade isopropanol (Sigma Aldrich) followed by drying with a filtered air stream. Silicon gaskets (Grace Bio-Labs) to hold the sample drops were cleaned in the same procedure immediately prior to measurement. All mass photometry measurements were performed at room temperature using DPBS without calcium and magnesium (ThermoFisher). The instrument was calibrated using a protein standard mixture: β-amylase (Sigma-Aldrich, 56, 112 and 224 kDa), and thyroglobulin (Sigma-Aldrich, 670 kDa). Before each measurement, 15 µL of DPBS buffer was placed in the well to find focus. The focus position was searched and locked using the default droplet-dilution autofocus function after which 5 µL of protein was added to make final concentration at 15 nM and pipetted up and down to briefly mix before movie acquisition was promptly started. Movies were acquired for 60 s (3000 frames) using AcquireMP (version 2.3.0; Refeyn Ltd) using standard settings. All movies were processed and analyzed using DiscoverMP (version 2.3.0; Refeyn Ltd).

### Biolayer interferometry (BLI)

Affinity determination measurements were performed on the Octet RED96 (Sartorius). All assays were performed using streptavidin (SA) coated biosensors (Sartorius) in kinetics buffer (PBS pH 7.4, 0.5 mg/mL BSA, and 0.01% (v/v) Tween-20) at 25°C. Biosensors were equilibrated for 30 min prior to beginning the assay. Assay step order and corresponding times were as follows: equilibration (30 s), loading (90 s), baseline (30 s), association (10 s), and dissociation (15 s). Biotinylated FAP and DPP4 (2 µg/mL, AcroBiosystems) were loaded onto SA sensors to a response of 0.5 nm. Association measurements were performed using a dilution series of MBP-I3 from 0.25 to 10 µM. Baseline drift was corrected by subtracting the response of a ligand-loaded sensor in kinetics buffer. Data analysis was performed with Octet Data Analysis 11.1 software using a global fit 1:1 model to determine affinity and kinetic parameters. Affinity and kinetic data reported are representative of three independent experiments.

### Cryo-EM sample preparation and data acquisition

The cross-linked FAP with SUMO-I3 sample (0.4 mg/mL) was vitrified on QuantiFoil R1.2/1.3 300 mesh copper grids (SPT Labtech) in 20 mM Tris-HCl pH 8.0 and 150 mM NaCl using a Vitrobot Mark IV (ThermoFisher). Grids were glow discharged for 60 s, –15 mA on a PELCO easiGlow (Ted Pella) system. Sample (3 µL) was applied to grids in the Vitrobot chamber (4 °C and 95% humidity) and blotted for three seconds with −5 blotting force before plunge-freezing in liquid ethane. Data were collected on a Titan Krios G3 microscope (300 kV) using SerialEM with a K3 direct electron detector (Gatan). A total of 5,450 movies were collected at a pixel size of 0.43 Å/pixel (super-resolution mode) with a dose of ∼65 electrons/A^2^, exposure time of 2.43 seconds, 45 frames, and a defocus range of −0.5 to −2.5 µm.

### Cryo-EM image processing and 3D reconstruction

Movies were subject to patch motion correction and patch CTF estimation in cryoSPARC^45^. Initial particle picks were performed on a subset of data using the blob picker followed by two-dimensional (2D) classification to generate 2D templates for template-based picking on the full dataset in cryoSPARC Live. Particles (2,293,048) were extracted using a 300-pixel box size fourier cropped to 150 pixels (1.72 Å/pixel) and cleaned with multiple rounds of 2D classification. The volume from streaming refinement in cryoSPARC Live was fed into heterogeneous refinement in cryoSPARC using 3 classes to isolate classes of FAP + SUMO-I3 and FAP + 2 SUMO-I3. Two classes from this Hetero refinement job were selected to re-extract separate particle stacks for FAP + SUMO-I3 (696,327) and FAP + 2 SUMO-I3 (356,520). Particles were extracted using a 300-pixel box size (0.86 Å/pixel) and used separately for non-uniform refinement in either C1 or C2 symmetry. A 3D classification job was used to further clean each particle stack, which resulted in final particle numbers of 272,711 for FAP + SUMO-I3 (C1 symmetry) and 238,246 for FAP + 2 SUMO-I3 (C2 symmetry). Processing was done initially in cryoSPARC version 3.1 and finished in version 3.3.1. Final maps were post-processed using DeepEMhancer and used for visualization.

### Model building and refinement

An initial model for complex of FAP with SUMO-I3 sdAb was generated by the crystal structure (PDB 1Z68) and Alphafold^46^. The model was initially docked into the density map using Fit in Map in Chimera^47^. Manual model building was performed in Coot^48^ and refinement using real-space refinement in Phenix^49^. Figures were generated in Chimera and PyMOL. Software used for data processing, model building, and refinement except for cryoSPARC were curated by SBGrid^50^.

### In silico rational design

The cryo-EM structure of FAP-I3 was imported into the Bioluminate package (Schrodinger Release 2023-3) and passed through the Protein Preparation Wizard. The FAP-I3 complex interface was manually inspected to come up with rational mutations that could both enhance and disturb the complex. Positions V107 and S109 in I3 were chosen as sites that could create additional interactions with FAP. Next, the Residue Scanning Module in Bioluminate was used to introduce mutations into I3. Stability and affinity calculations were performed optimizing for the affinity, and backbone minimization was used with a cutoff of 5 Å. Interface interactions were determined using the Protein Interaction Analysis Module.

---

**Figure S1.**
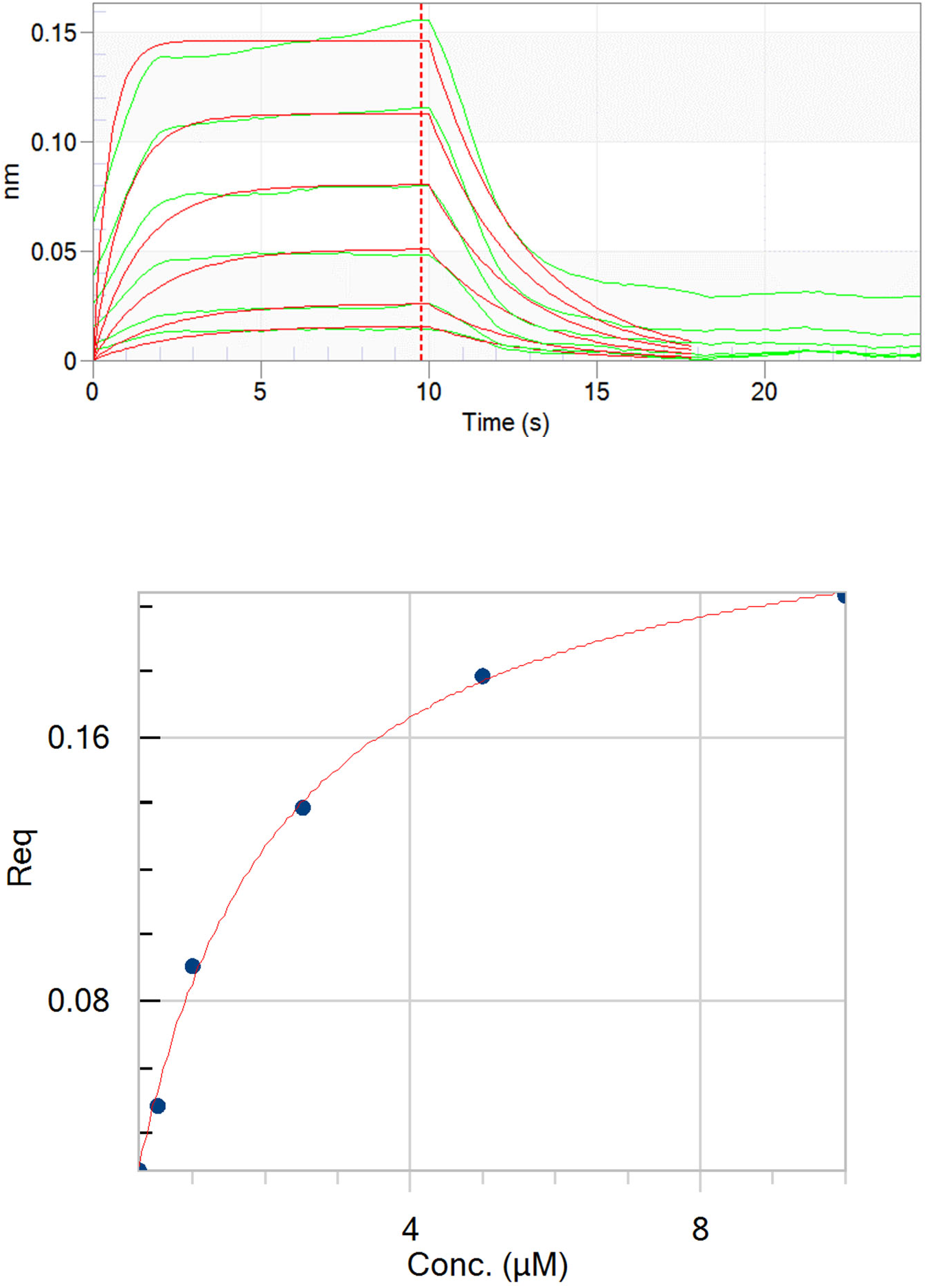
Binding affinity of FAP-MBP-I3 by Biolayer Interferometry (BLI) (A) Kinetic fits (red lines) for 1:1 model from FAP-MBP-I3 binding data (green lines). R^2^ = 0.98 X^2^ = 0.01 (B) Steady state analysis of FAP-MBP-I3 binding data. R^2^ = 0.998 R_max_ = 0.2403 ± 0.004. K_d_ = 1.80 ± 0.09 µM

**Figure S2.**
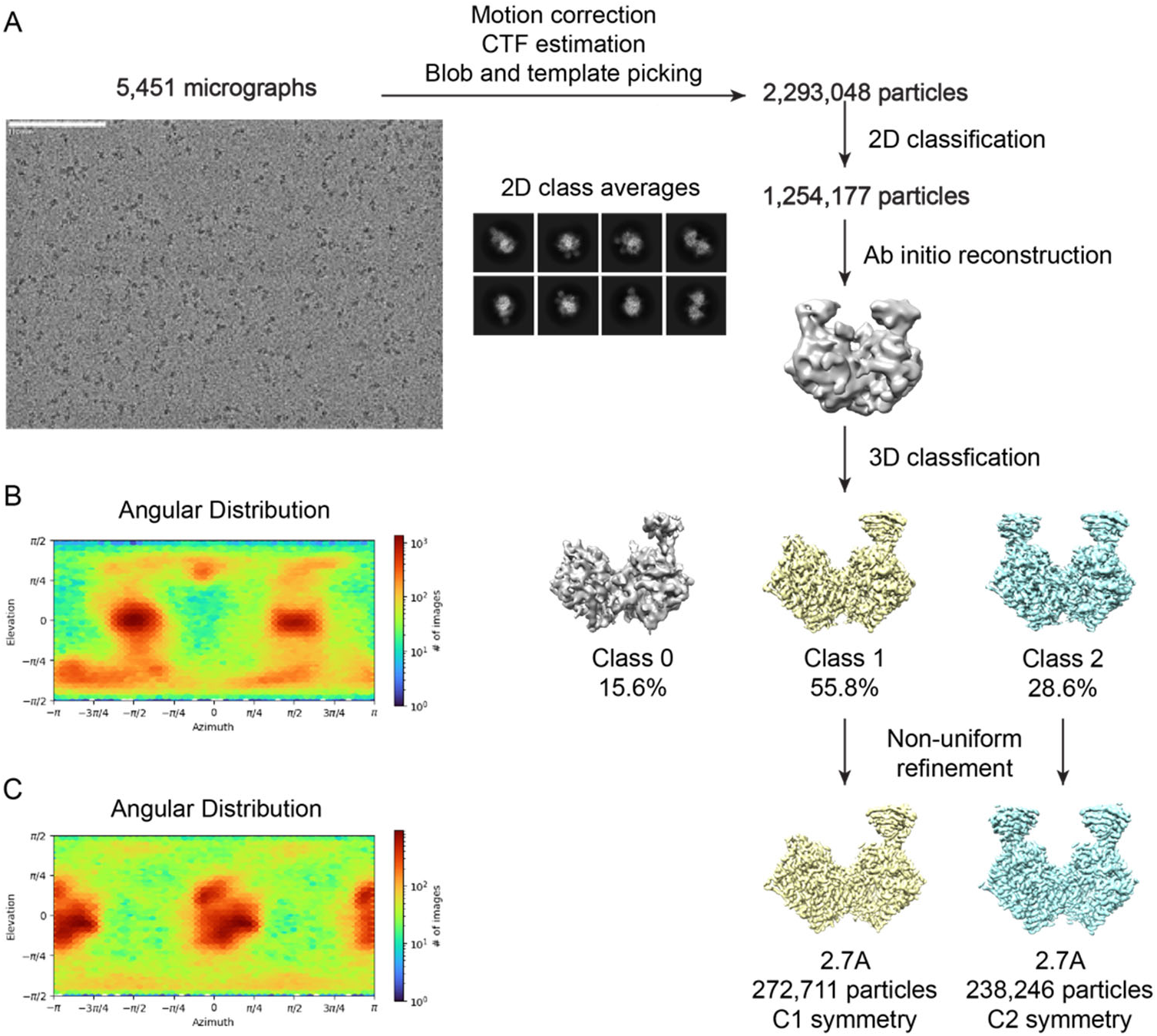
Cryo-EM data processing workflow. (A) Processing workflow used in cryoSPARC to obtain both reconstructions of one and two SUMO-I3 molecules bound to FAP. (B) Particle view distribution for one SUMO-I3 bound to FAP (C1 symmetry). (C) Particle view distribution for two SUMO-I3 bound to FAP (C2 symmetry).

**Figure S3.**
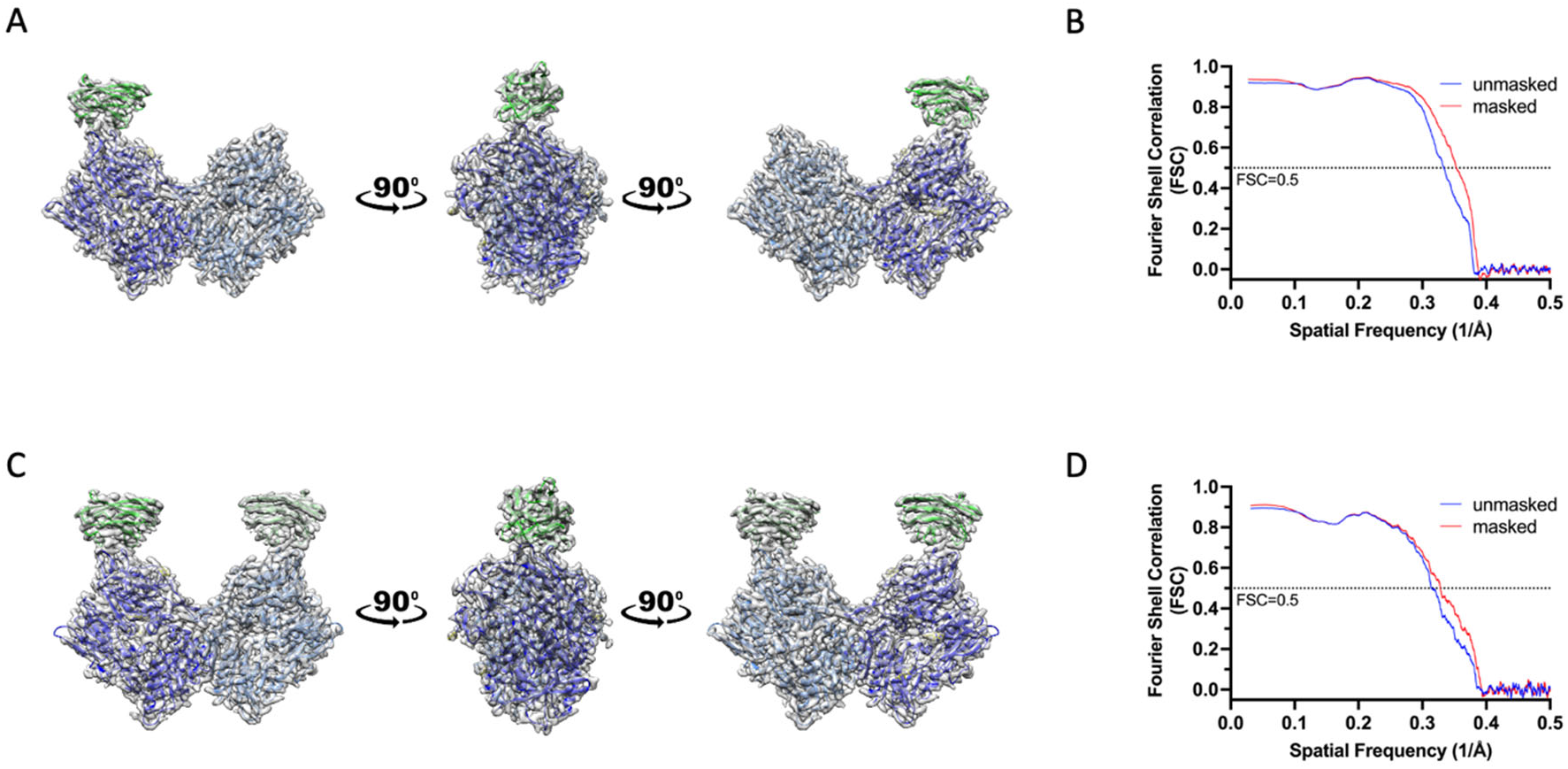
Map and model overlay. (A) Cartoon model of FAP-I3 overlaid with the final reconstruction map showing various orientations. (B) Map-to-model FSC for FAP-I3. (C) Cartoon model of FAP + 2 SUMO-I3 overlaid with the final reconstruction map showing different orientations. (D) Map-to-model FSC for FAP + 2 SUMO-I3.

**Figure S4.**
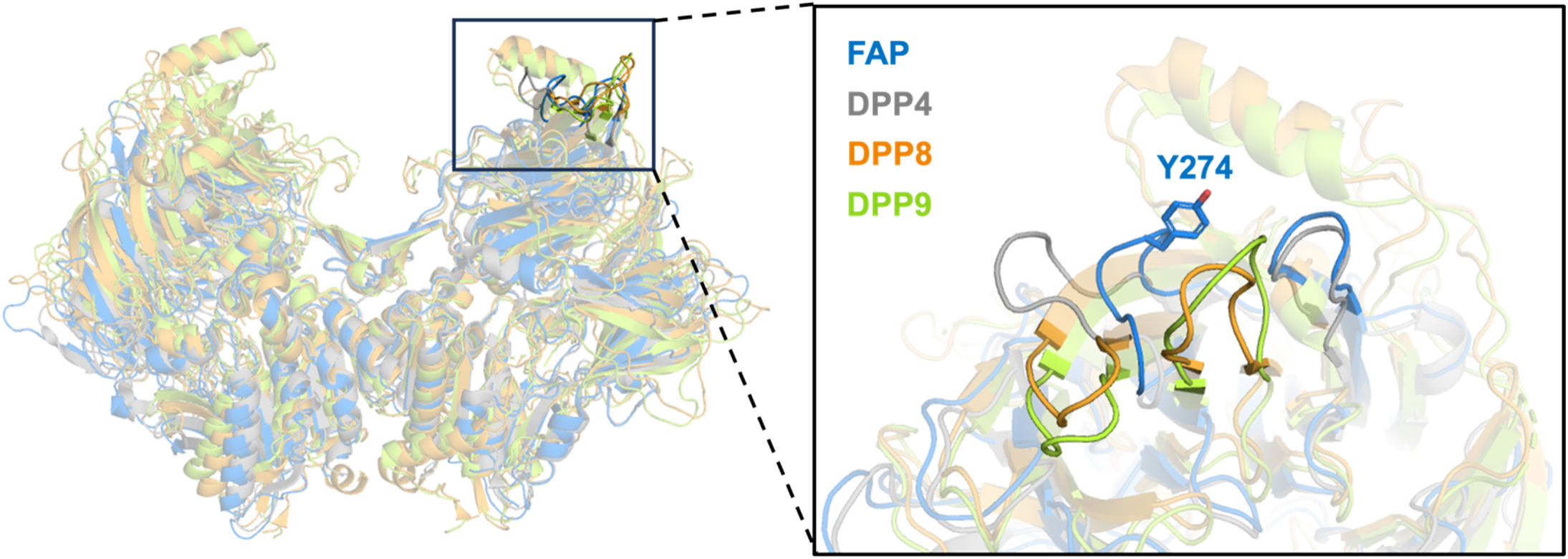
Overall alignment of FAP with DPPs and comparison of I3 epitope region of FAP to structurally aligned regions in DPPs. PDB codes: FAP (1Z68), DPP4 (2ONC), DPP8 (6EOO), DPP9 (7A3F).

## Acknowledgements

Dr. Wadas gratefully acknowledges the NCI (R21-CA219899 and R21-CA227709), the DoD (W81XWH-19-1-0046), The Department of Radiology and The Holden Comprehensive Cancer Center Gift Fund for research support. The authors acknowledge use of the University of Iowa Central Microscopy Research Facility, a core resource supported by the University of Iowa Vice President for Research, and the Carver College of Medicine. The Octet system, which is located within the Protein and Crystallography Facility, is supported by The Holden Comprehensive Cancer Center at The University of Iowa and its National Cancer Institute Award P30CA086862. Preliminary cryo-EM data was collected at the Iowa State University Cryo-EM facility on the Glacios 200kV TEM and Gatan K3 detector. We thank Dr. Puneet Juneja for his assistance in grid screening and initial data collection. The authors would also like to acknowledge Dr. Stefan Steimle from the Beckman Center for Cryo Electron Microscopy at the University of Pennsylvania Perelman School of Medicine for support with final cryo-EM data collection (RRID: SCR_022375). Finally, the authors acknowledge Dr. Ernesto Fuentes, Ph.D. for his insightful comments and fruitful discussions during the preparation of this manuscript.

## Author Contributions

The author contributions are as follows: Conceptualization, T.W. and N.J.S.; methodology, D.N.P., Z.X., and A.S.; data curation, D.N.P., Z.X., and A.S.; writing—original draft preparation, T.W., Z.X., and N.J.S. writing— review and editing, T.W., and N.J.S.; supervision, T.W., and N.J.S.; project administration, T.W.; funding acquisition, T.W. All authors have read and agreed to the published version of the manuscript.

## Materials and Correspondence

Correspondence and requests for materials should be addressed to Thaddeus J. Wadas or Nicholas J. Schnicker.

